# A NMF-based approach to discover overlooked differentially expressed gene regions from single-cell RNA-seq data

**DOI:** 10.1101/543447

**Authors:** Hirotaka Matsumoto, Tetsutaro Hayashi, Haruka Ozaki, Koki Tsuyuzaki, Mana Umeda, Tsuyoshi Iida, Masaya Nakamura, Hideyuki Okano, Itoshi Nikaido

## Abstract

Single-cell RNA sequencing has enabled researchers to quantify the transcriptomes of individual cells, infer cell types, and investigate differential expression among cell types, which will lead to a better understanding of the regulatory mechanisms of cell states. Transcript diversity caused by phenomena such as aberrant splicing events have been revealed, and differential expression of previously unannotated transcripts might be overlooked by annotation-based analyses.

Accordingly, we have developed an approach to discover overlooked differentially expressed (DE) gene regions that complements annotation-based methods. We applied our algorithm to two datasets and discovered several intriguing DE transcripts, including a transcript related to the modulation of neural stem/progenitor cell differentiation.

## Background

The advancement of single-cell technology has enabled to investigate various tissues [1, 2] and species [3, 4] with single-cell RNA sequencing (scRNA-seq), which enables comprehensive cell typing and the elucidation of cell compositions and dynamics. In particular, scRNA-seq can reveal the subtle differences among cell states, such as intermediate stages of differentiation. By investigating differentially expressed (DE) genes among such cell states, we can elucidate regulatory processes including cell fate determination [5]. In addition to traditional gene-level differential expression analyses, various novel analyses have been proposed for scRNA-seq studies, including the detection of differential distributions of expression levels [6] and differential splicing [7, 8], isoform-level differential pattern analysis [9], discriminative learning approach for differential expression analysis [10], and dynamic prediction through the comparison of spliced and unspliced mRNAs [11]. Thus, the development of various computational analysis methods that utilize information at the single-cell level is essential to advance the current understanding of RNA biology.

Recent comprehensive analyses of RNA-seq data have revealed the existence of various overlooked transcripts. For example, a comprehensive tumor analysis revealed that many tumors contain aberrant splicing patterns (neojunctions) that are not detected in normal samples [12]. Additionally, numerous genetic variants are related to aberrant splicing associated with certain diseases [13]. Therefore, it is important to detect novel splicing patterns, as well as detect differential expression of annotated transcripts. The transcriptomes of unstudied cell types, including rare cell types, can be revealed by scRNA-seq analyses, and we can now discover such cell type-specific splicing events.

In addition to major types of alternative splicing (AS), underappreciated classes of AS events, such as retained introns and microexons, are known to have essential roles, for example, in neuronal development [14]. Intron retention, which is common in tumors, can generate peptides and be a source of neoepitopes for cancer vaccines, and therefore the detection of novel intron retention events is medically important [15]. Furthermore, alternative polyadenylation, which produces isoforms that have 3’-untranslated regions (UTRs) of different lengths, is also known to be associated with several biological processes [16].

To reveal such complex AS patterns, several computational approaches have been developed that can detect previously unannotated splicing patterns. For example, spliced aligned reads (exon–exon junction reads) are beneficial in identifying the spliced mRNA structures [17, 18]. As another example, non-negative matrix factorization (NMF) has been used to decompose data into essential patterns and predict AS patterns from microarray data [19] and RNA-seq data [20].

In addition to these complex AS patterns, other types of transcripts, such as antisense transcripts transcribed from gene regions, are known to be essential regulators of gene expression [21]. In light of such complex transcript structures, typical differential expression analysis based on previously annotated transcript structures might overlook some important DE genes. To find DE genes without relying on existing annotation, distinct approaches have been proposed that identify DE regions from read coverage data [22, 23].

In single-cell technologies, full-length scRNA-seq data such as Smart-Seq [24, 25] provide powerful data that can reveal these complex transcript structures. Other scRNA-seq protocols, such as SUPeR-Seq [26], which can capture non-poly(A) transcripts, will also be useful to detect various overlooked DE transcripts. In particular, we have developed a single-cell full-length total RNA-seq (RamDA-seq) method and have validated that it precisely captures full-length transcripts and also captures various types of RNAs such as enhancer RNAs [27]. By utilizing such scRNA-seq data, we can perform differential expression analyses between cell states more precisely.

Accordingly, we have developed an approach to discover **O**verlooked **D**ifferentially **E**xpressed **G**ene **R**egions (ODEGRs), which is derived from several kinds of transcripts such as novel AS patterns, intron retention, and antisense transcripts, to complement the annotation-based differential expression analysis of single-cell data (Fig.1). Our approach utilizes the composition of scRNA-seq data, which contain information from many samples (i.e., cells), and decomposes the mapped count data for gene regions using NMF. With NMF, we can computationally extract reproducible signals corresponding to transcript structures and their associated expression profiles without relying on transcript annotations (Fig.2(a)). In addition, the non-negative constraint of NMF, which is its principal difference from other matrix decomposition methods, is effective in preserving the relation of the magnitude of expression. Next, we developed the following scores for a gene region: 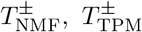 and Δ*T*_NMF–TPM_. 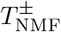 represents the scores that quantify the differential expression levels between two groups based on the NMF result (Fig.2(b)), while 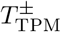 represents the scores that quantify the differential expression levels for annotation-based expression data (Fig.2(c)). Thus, Δ*T*_NMF–TPM_ represents the score that quantifies the differential expression that is not detected in the annotation-based approach (Fig.2(d)). We investigated gene regions with high Δ*T*_NMF–TPM_ values in order to discover ODEGRs.

**Figure 1.**
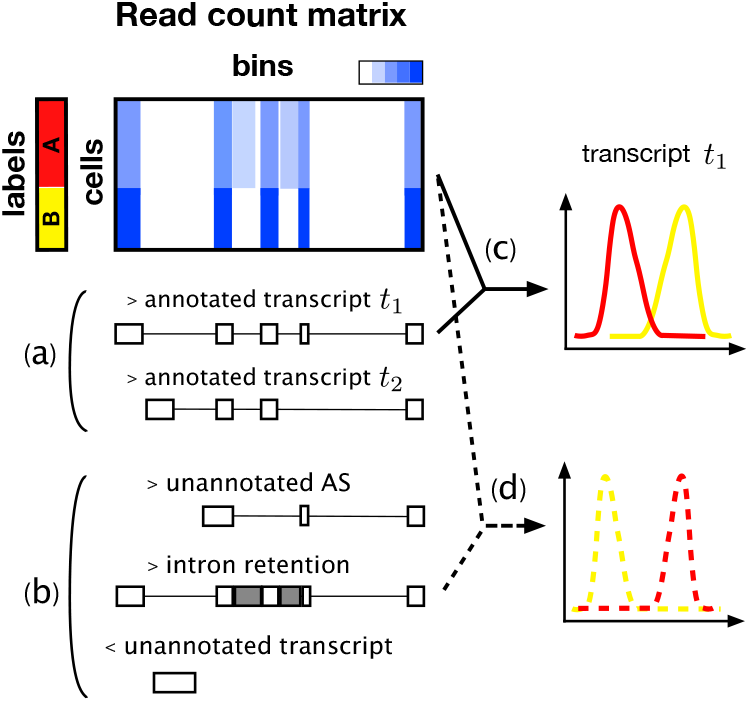
Graphical abstract of the overlooked differentially expressed gene region (ODEGR). Coverage of scRNA-seq data and annotated transcripts in the region (a) and previously unannotated transcripts such as novel alternative splicing patterns, intron retention, and unannotated antisense transcripts (b). Although annotation-based expression profiling and the following differential expression analysis is an effective approach to find DE transcripts (c), such a method might overlook the differential expression of unannotated transcripts (d).

**Figure 2.**
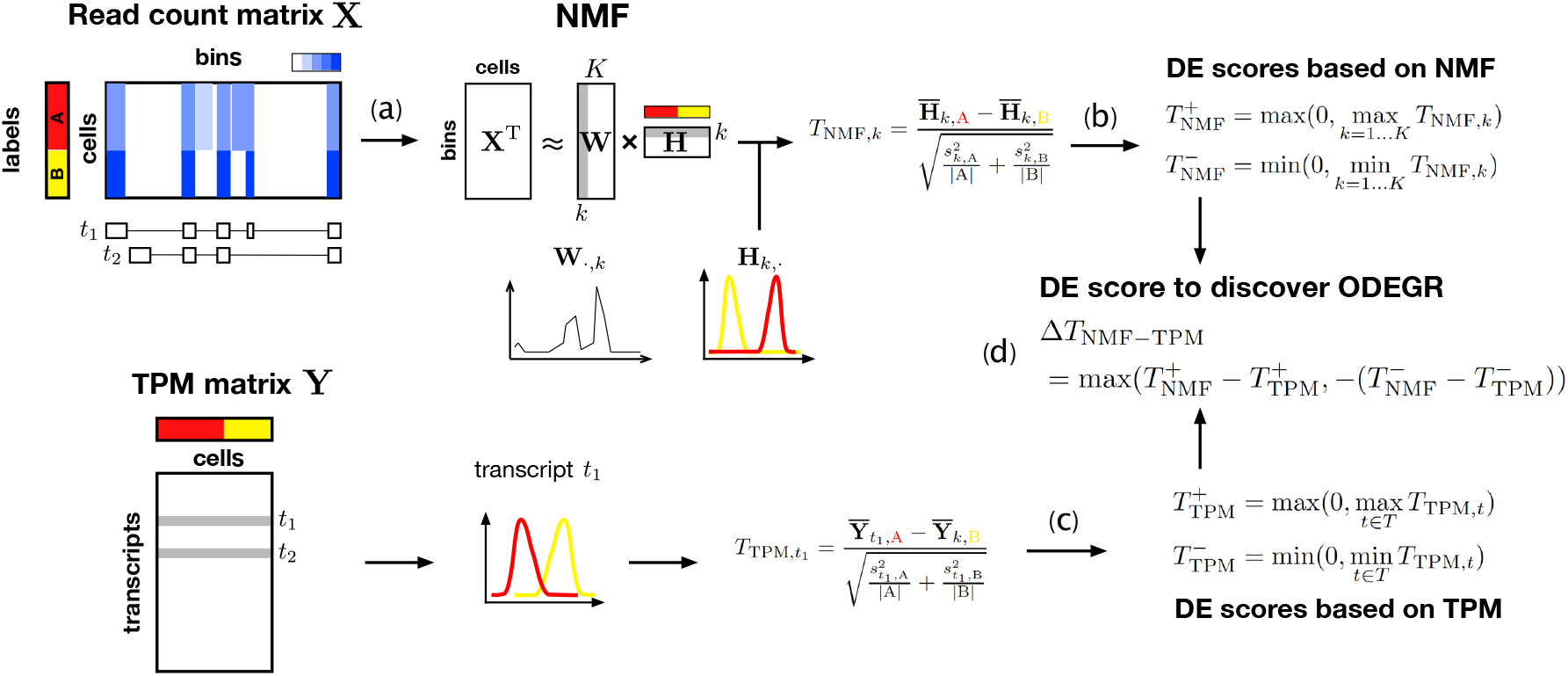
Graphical abstract of our algorithm to discover ODEGRs. First, we use non-negative matrix factorization (NMF) to decompose the mapped read count matrix (**X**) for a gene region (a), and then use *t*-statistics to quantify the differential expression level while keeping the positive maximum and negative minimum values (b). We also quantify the differential expression level using an annotation-based expression profile (in a transcripts per million (TPM) matrix) (c). Finally, we quantify the unexpectedness of differential expression based on the above values (d).

We applied our algorithm to two real datasets: (1) mouse embryonic stem (ES) cells and primitive endoderm (PrE) cells and (2) neural stem cells (NSCs) derived from human induced pluripotent stem (iPS) cells. First, we evaluated whether the NMF-based approach could quantify and find local DE regions from simulated data. We also evaluated whether it could detect AS switches within a gene, as determined by annotation-based analysis. Our algorithm was indeed able to detect such DE regions without relying on transcript annotations. Then, we applied our method to real datasets to detect ODEGRs and found several intriguing examples. From the perspective of previous research, our results correspond, for example, to unannotated splicing patterns, antisense transcript, and unannotated 3*’*-UTRs of adjacent genes. In particular, some ODEGRs are related to critical regulatory mechanisms such as the modulation of differentiation and tissue-specific imprinting. Thus, our novel differential expression analysis method identified some important ODEGRs and can complement annotation-based methods, making it a useful method for analysis in the increasing number of scRNA-seq experiments.

## Results

### NMF-based approach for discovering ODEGR

In this research, we focused on detecting DE gene regions that were overlooked in the differential expression analysis of previously annotated transcripts from mapped read count data. We divided a gene region into 100-bp bins and described a read count matrix for a gene region with a *C* × *L* matrix **X**, where *C* is the number of cells and *L* is the number of bins. First, we decomposed **X** into two non-negative matrices (using non-negative matrix factorization):

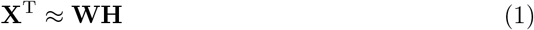

where **W** and **H** are *L* × *K* and *K* × *C* non-negative matrices (*K* is the factorization rank) referred to as “metagenes” and “metagene expression profiles” in previous studies, respectively [28, 29]. In this research, we hypothesized that **W** corresponds to the transcript structure including splicing patterns and that **H** corresponds to the expression for each structure in each cell.

Second, we quantified the differential expression level of a structure *k* ∈ (1…*K*) between two groups *A* and *B* based on Welch’s *t*-test:

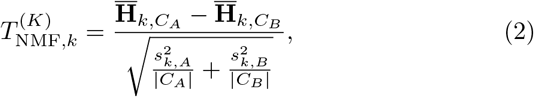

where *C*_*A*_ is the list of cells whose labels are *A* and 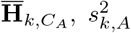 and |*C*_*A*_| are the sample mean of **H**_*k,·*_, variance, and size of group *A*, respectively. Owing to the non-negative constraint, the relation between the two groups (i.e., 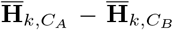 can be greater or smaller than 0) will be consistent with the relation in the original expression space. Our goal was to identify overlooked differential expression, and therefore, such relations, as well as their absolute values, were effective indicators for discovering ODEGRs. Therefore, we defined the following two scores, which correspond to the relation 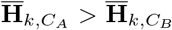 and 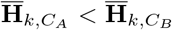, respectively:

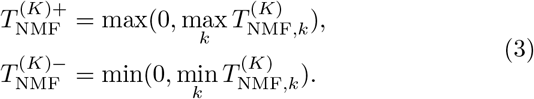

In NMF, the factorization rank (*K*) must be decided in advance, and the value is critical for analytical results. The various transcript structures cannot be separated with small *K* values and are excessively separated with large *K* values. In either case, the expression profiles become ambiguous, and we might overlook the DE regions if an inappropriate *K* value is selected. Therefore, we decomposed the data with several *K* values (*K* ∈ (2, 5, 10) in this research) and calculated the positive maximum and negative minimum values:

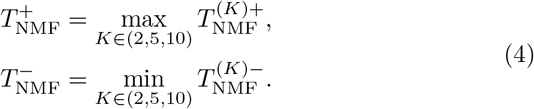

Next, we defined similar scores for the TPM (transcripts per million) matrix, which represents the expression profile based on annotated transcripts (we used log_10_(TPM + 1) in actuality). We described the list of transcripts for the gene region using *T* and calculated Welch’s *t*-statistic as before for a transcript *t* ∈ *T,* which is referred to as *T*_TPM,*t*_. Then, the scores for the gene region were defined by the positive maximum and negative minimum among transcripts of the gene as follows:

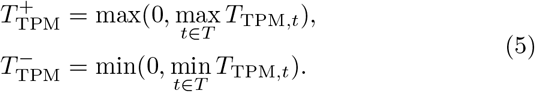

Lastly, we developed a score to detect ODEGRs as follows:

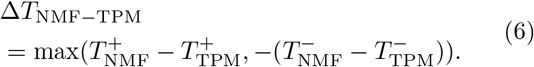

Because these is no global NMF optimization algorithm, we calculated Δ*T*_NMF–TPM_ using three different random seeds and also used minimum Δ*T*_NMF–TPM_ to obtain reliable ODEGRs (see the Methods section).

We also developed a score Δ*T*_NMF–Mean_ that measured the overlooked differential expression merely using the mean of the coverage. We used this score to evaluate whether the NMF-based approach separates the signal and detects complex DE patterns. We calculated the mean of the logarithm of data for a cell *c* 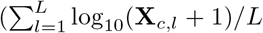, where *L* is the number of bins) as well as the corresponding Welch’s *t*-statistic as before and Δ*T*_NMF–Mean_ likewise.

### Dataset

In this research, we used scRNA-seq data from the following two experiments.

#### mES-PrE dataset

The first dataset is derived from mouse ES cells and primitive endoderm (PrE) cells subjected to RamDA-seq and was examined in our previous study [27]. We used the data from 5G6GR mouse ES cells samples at 0 and 72 h after dexamethasone induction and defined the cell type at each time point as ES cells (92 cells) and PrE cells (93 cells), respectively.

#### hNSC-NC dataset

The second dataset corresponds to human neural stem cells (NSCs) derived from iPS cells measured by RamDA-seq. There is heterogeneity within the population, and some subpopulations other than the NSC subpopulation were identified (Additional file 1: Fig. S1). After clustering these cells and defining the cell types based on marker gene expression, we identified 515 NSCs and 80 partially differentiated neural cells (NCs).

### Validation on simulation dataset

At first, we investigated the performance of NMF-based differential expression quantification and whether our approach can quantify the local differences in a region using simulation data. The simulation data were generated from the mES-PrE dataset such that the data matrix includes local DE patterns of length *L’* (see Methods section for detailed procedure). Then, we regarded the simulation and raw data as positive-control and negative-control datasets, respectively. We evaluated the ability to detect local DE regions based on Δ*T*_NMF–Mean_. We also compared the performance when we used *K* = (2, 5, 10), as mentioned in Eq. (4), or one fixed value (i.e., *K* = 2, 5, or 10) for calculating 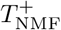 and 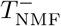.

The area under the ROC curve (AUROC) values for *all, K* = 2, *K* = 5, and *K* = 10 were 0.98, 0.93, 0.93, and 0.90, respectively for simulation data with *L*’ = 100 (Fig.3(a)). The AUROC values for the *L*’ = 50 dataset were 0.98, 0.91, 0.96, and 0.93, respectively, and those for the *L*’ = 10 dataset were 0.94, 0.64, 0.86, and 0.94, respectively (Fig.3(b),(c)). In all cases, our algorithm using multiple *K* values showed high performance, and therefore, our NMF-based approach is useful for discovering various local differences.

**Figure 3.**
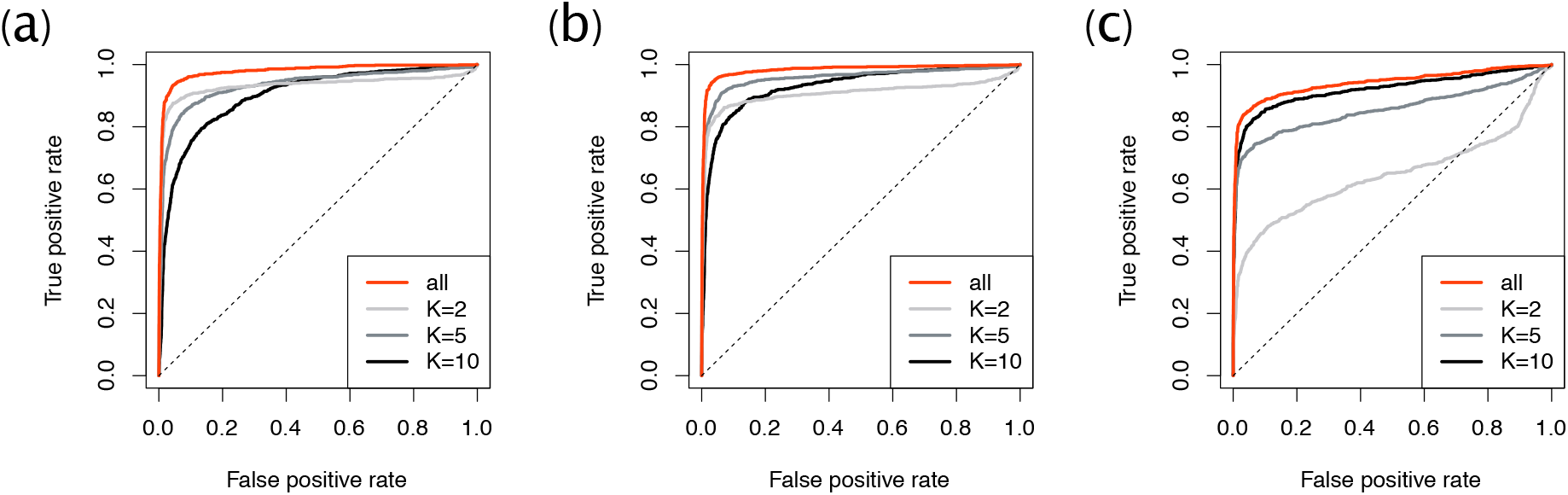
The ROC curves for the simulated dataset. Simulation results for (a) *L*’ = 100, (b) 50, and (c) 10, where *L*’ is the length of local differential expression patterns.

### Validation with alternative isoform expression

We also investigated whether the NMF-based approach can quantify the complex DE patterns associated with genes that have alternative isoform expression. Based on the TPM matrix calculated from the annotation, we defined the positive-control and negativecontrol datasets. The former consists of the gene set with different isoforms expressed in different groups, while the latter consists of the remaining genes (see the Methods section for detailed definitions). Then, we evaluated the ability to detect such complex DE patterns based on Δ*T*_NMF–Mean_.

The positive-control examples of alternative isoform expression in the mES-PrE dataset were *Frmd4a* and *Pde4d*, which are known for frequent transcription start site (TSS) switching events [30] (Fig.4(a),(b)). Based on our criteria, both *Frmd4a* and *Pde4d* were highly ranked (53rd and 23rd out of 4,965 genes, respectively).

**Figure 4.**
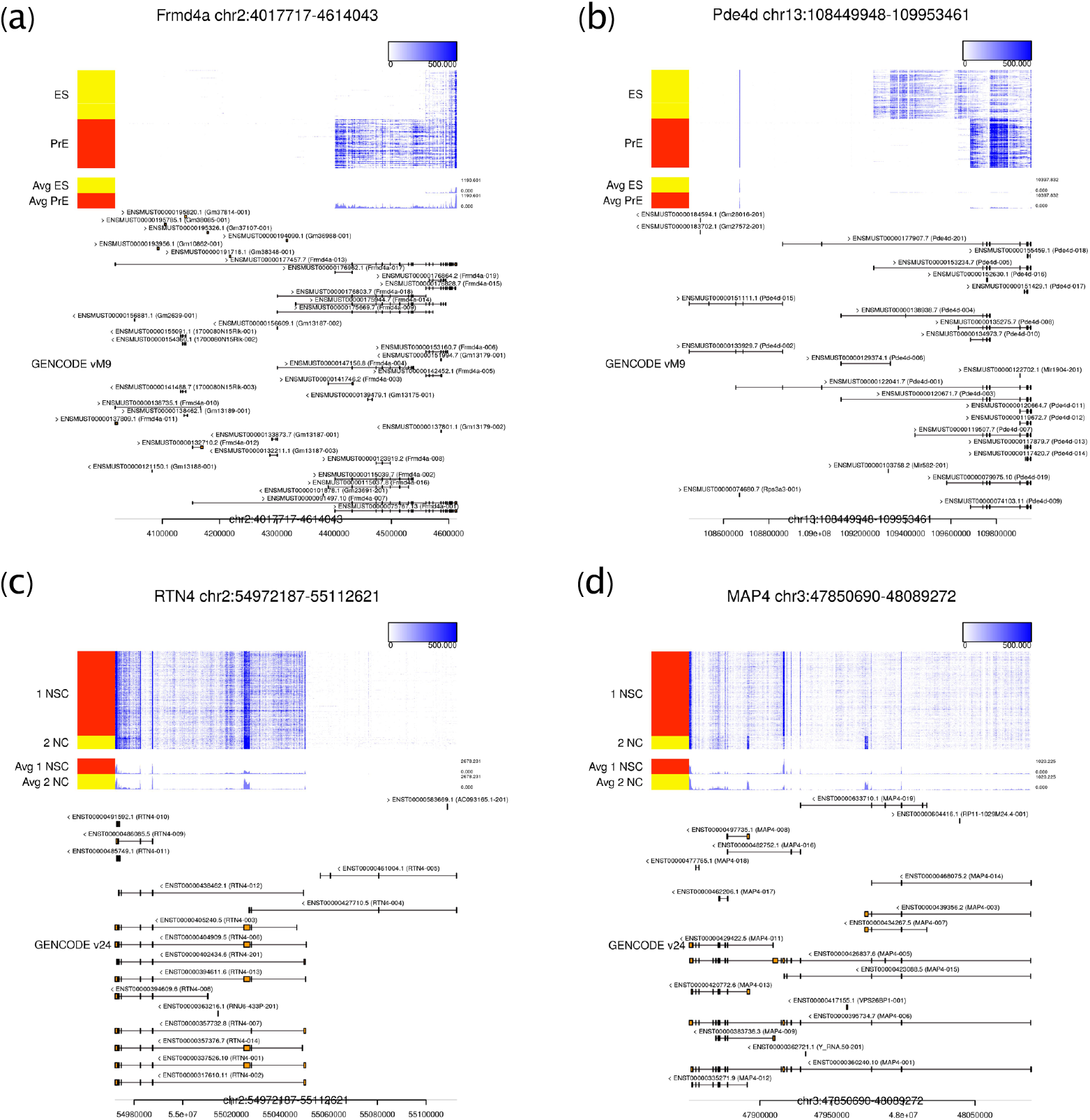
Examples of alternative isoform expression. The visualizations of read coverage and transcript annotations for (a) *Frmd4a*, (b) *Pde4d*, (c) *RTN4*, and (d) *MAP4*, respectively. (a) and (b) are the examples from the mES-PrE dataset, while (c) and (d) are the examples from the hNSC-NC dataset. These figures are visualized with Millefy, which provides genome-browser-like visualizations of scRNA-seq datasets https://github.com/yuifu/millefy.

The examples in the hNSC-NC dataset were *RTN4*, also known as *NOGO*, which encodes the Nogo-A iso-form that contains exon 3 and is expressed in neural precursor cells [31] (Fig.4(c)), and *MAP4*, which is known for its alternative isoform expression across neural cell types [32] (Fig.4(d)). These genes were highly ranked in our criteria (40th and 1st out of 6,491 genes, respectively.) Thus, the typical genes with alternative isoform expression are highly ranked in our criteria Δ*T*_NMF–Mean_.

Overall, the AUROC values (for threshold 15) were about 0.79 and 0.83 for the mES-PrE and hNSC-NC datasets, respectively (Fig.5). Although our algorithm overlooked some alternative expression patterns, the high AUCROC values demonstrated the effectiveness of our algorithm for discovering previously unannotated DE transcripts.

**Figure 5.**
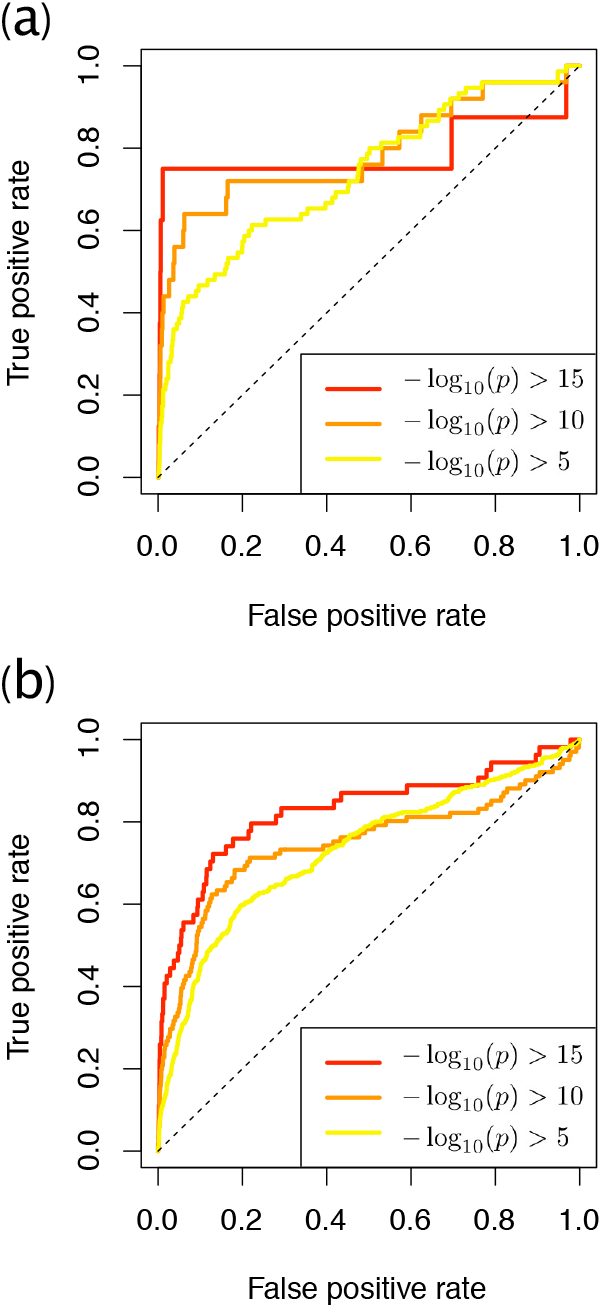
The ROC curves for detecting genes with alternative isoform expression. The results for the (a) mES-PrE and (b) hNSC-NC datasets.

### Discovery of ODEGRs

Next, we investigated the existence of ODEGRs by using Δ*T*_NMF_ – _TPM_. In brief, the values of Welch’s *t*-statistics based on NMF (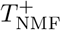 and 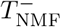) and TPM (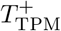 and 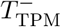) were highly correlated (Pearson’s correlation coefficients for the mES-PrE dataset and hNSC-NC dataset were about 0.83 and 0.84, respectively), and large Δ*T*_NMF–TPM_ values were observed for only a small fraction of genes (Additional file 1: Fig. S2). Therefore, we ranked genes by Δ*T*_NMF–TPM_ in descending order to identify ODEGRs. (The actual procedure of Δ*T*_NMF–TPM_ calculation is described in the Methods section.) Only a small fraction of genes had large positive values of Δ*T*_NMF–TPM_ (Additional file 1). Five genes in the mES-PrE dataset had Δ*T*_NMF–TPM_ *Z*-scores over 3 while 39 genes in the hNSC-NC dataset did. Although the number of ODEGRs discovered by our algorithm were few, several intriguing ODEGRs were identified.

#### mES-PrE Dataset

The read coverage and transcript annotation for the six highest-ranking genes in the mES-PrE dataset are shown in Fig.6.

**Figure 6.**
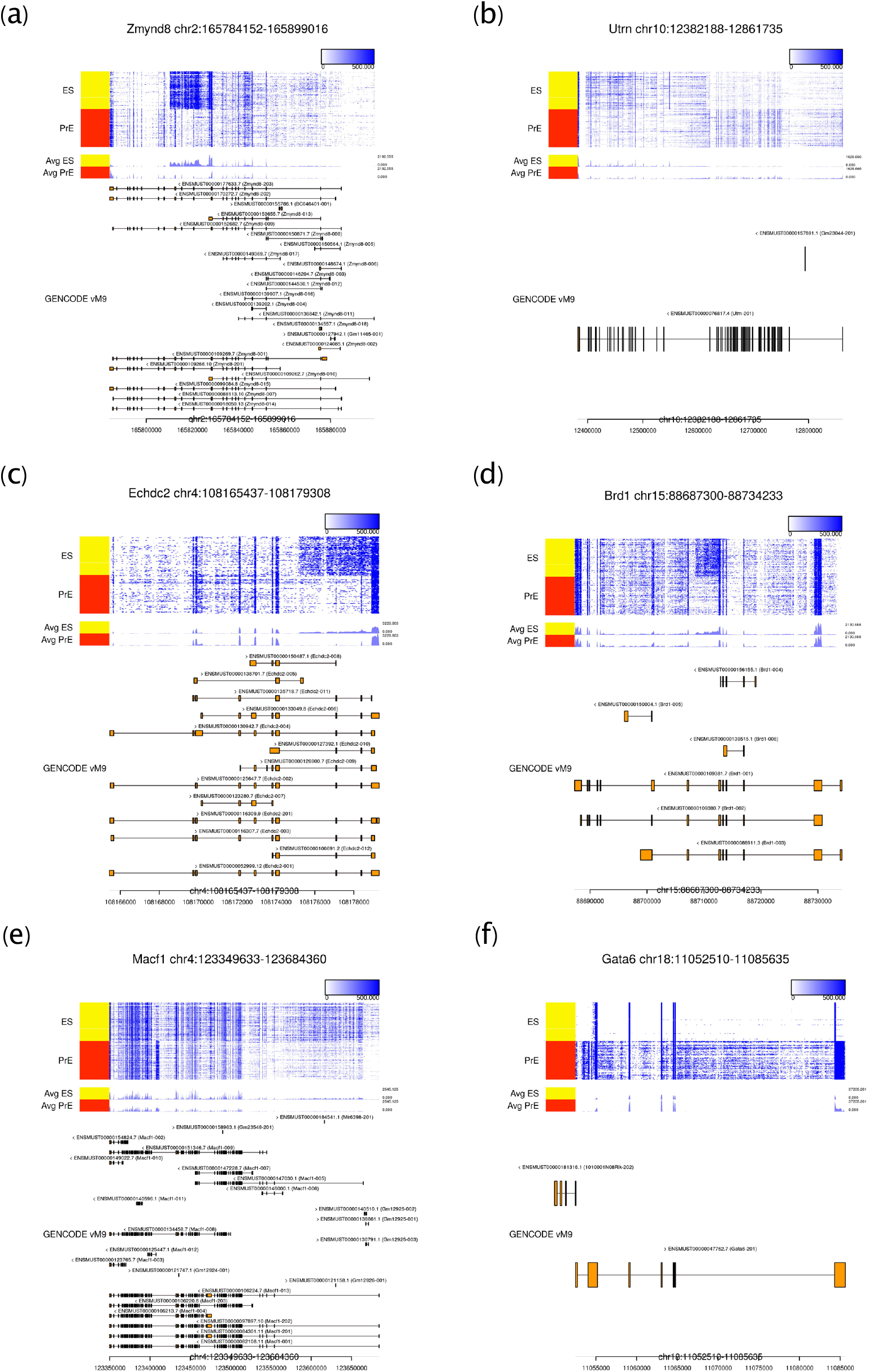
Examples of high-ranking genes in the mES-PrE dataset. The results for the six top-ranked genes (in descending order) (a) *Zmynd8*, (b) *Utrn*, (c) *Echdc2*, (d) *Brd1*, (e) *Macf1*, and (f) *Gata6* are visualized.

The 1st and 4th ranked genes were *Zmynd8* and *Brd1*, and numerous reads were mapped to the specific intron regions of these genes (Fig.6(a)(d)). The novel enhancer-associated antisense transcripts for these genes have previously been reported in mESCs [33], and this suggests that our approach can detect several kinds of DE transcripts, including antisense transcripts.

The 2nd ranked gene was *Utrn*, and two distinct coverage patterns of peaks that correspond to exons were observed in ES and PrE cells, respectively (Fig.6(b)). Since the annotation contains only one isoform, this DE pattern was overlooked in the annotation-based approach. We used GENCODE vM9 such that the analytical results were consistent with previous work [27], and we also considered the possibility that the latest annotation includes the isoforms corresponding to such patterns. We recalculated the TPM values using GENCODE vM18, and 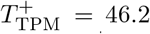 and 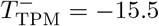 for vM18, in comparison to 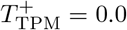 and 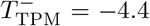 for vM9 (Additional file 1: Section 3.2 and Fig. S4). This result suggests the existence of DE transcripts that were not annotated in vM9. A similar result was observed for the 7th ranked gene *Arid5b* (Fig. S4). These results demonstrate the potential of our approach for discovering previously unannotated isoforms.

The 3rd ranked gene was *Echdc2*, which had numerous reads mapped to its 3*’* intron region (Fig.6(c)). Although such a pattern is consistent with intron retention, this mapping pattern is continued from adjacent gene *Zyg11a*, and the coverage at the 3*’* intron of *Echdc2* is correlated with coverage at the *Zyg11a* region (Additional file 1: Section 3.3 and Fig. S5). These results suggest that an unannotated long isoform of *Zyg11a* exists and overlaps with the *Echdc2* region.

The 5th ranked gene was *Macf1*, and numerous reads were mapped to the specific intron region of the gene in PrE cells (Fig.6(e)). An exon was annotated for the region in vM18, and the DE transcript including the exon was overlooked in differential expression analysis using vM9, which was also the case for *Utrn* and *Arid5b* (Fig. S4).

The 6th ranked gene was *Gata6* (Fig.6(f)). The exogenous *Gata6*, which lacks a 3*’*UTR end, is arbitrarily expressed in these ES cells. After dexamethasone induction, Gata6 is transported into the nucleus, ES cells differentiate into PrE cells, and the level of expressed endogenous *Gata6* increases. Because the annotation file does not include exogenous structure, annotation-based TPM cannot reflect the exogenous expression patterns, which resulted in high Δ*T*_NMF–TPM_ values.

#### hNSC-NC Dataset

In comparison to the results of the mES-PrE dataset, the results of the hNSC-NC dataset contained uninteresting patterns among the most highly ranked genes (Additional file 1: Section 3.1 and Fig. S3). Therefore, we show six high-ranking genes of great interest in the hNSC-NC dataset (Fig.7).

**Figure 7.**
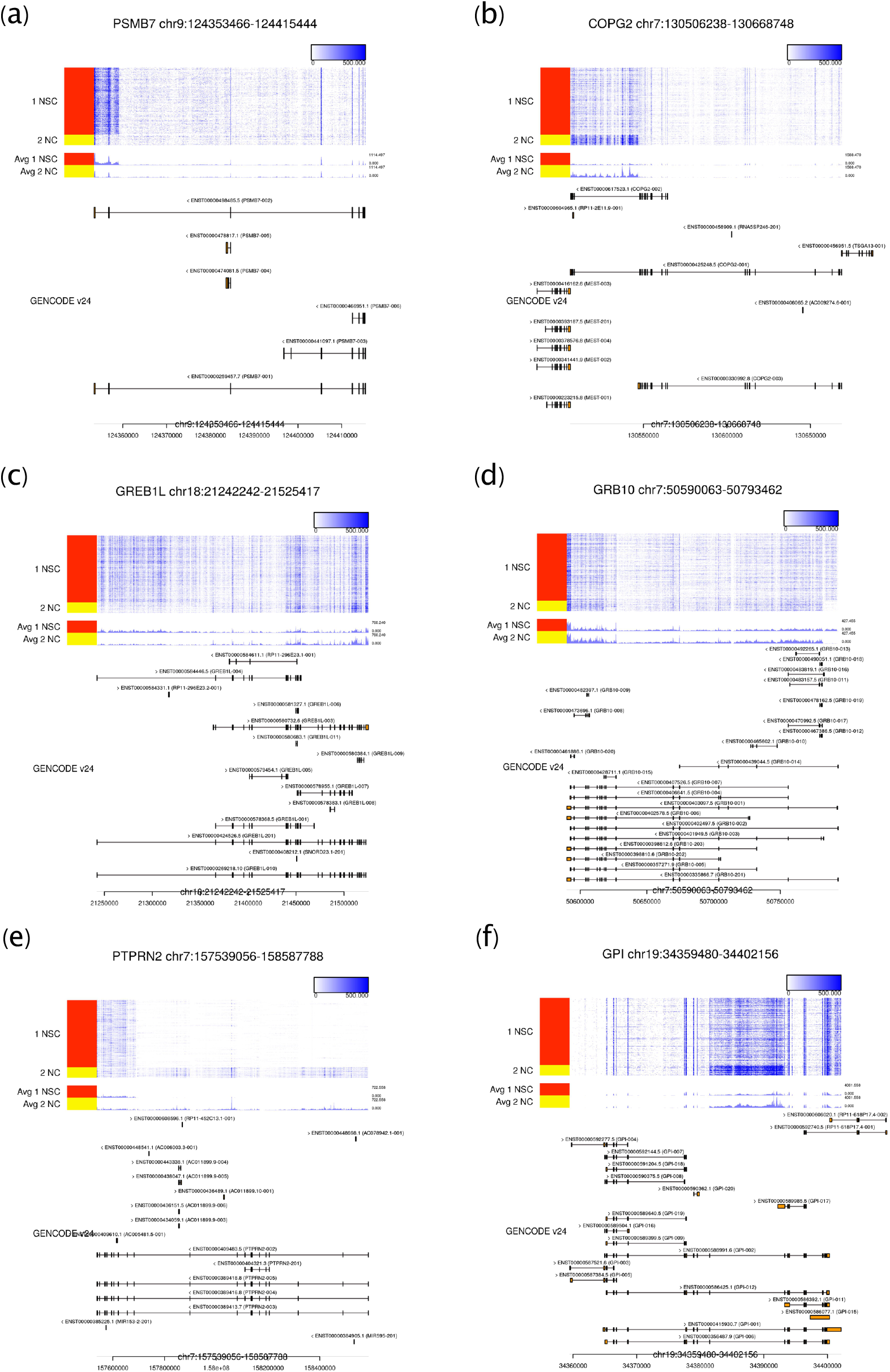
Examples of high-ranking genes in the hNSC-NC dataset. The results for (a) *PSMB7*, (b) *COPG2*, (c) *GREB1L*, (d) *GRB10*, (e) *PTPRN2*, and (f) *GPI*, the 2nd, 6th, 10th, 15th, 17th, and 18th ranked genes, respectively, are visualized.

The 2nd ranked gene was *PSMB7*, and many reads from NSCs were mapped to its 3*’* intron region, which is similar to the result for *Echdc2* in the mES-PrE dataset (Fig. 7(a)). The coverage pattern was continued from the adjacent gene *NEK6*, and the coverage of the intron region is correlated with the that of *NEK6* (Fig. S5). This result suggests the existence of an unannotated long transcript of *NEK6* that overlaps with the *PSMB7* region.

The 6th ranked gene was *COPG2*, and numerous reads were mapped to its 3*’* intron regions, resembling the results for *Echdc2* and *PSMB7* (Fig.7(b)). These reads are also likely to be derived from transcripts of the adjacent gene *MEST*, which may have an unannotated long transcript. Intriguingly, in mouse, *Mest* is an imprinted gene, and a long isoform of *Mest* (referred to as *MestXL*) is expressed in the developing central nervous system, which results in the repression of *Copg2* on the same paternal allele [34]. Therefore, the long transcript of *MEST* and the tissue-specific imprinting of *COPG2* depending on the long transcript are thought to occur in human. Thus, the detection of overlapping unannotated transcripts can be associated with regulatory mechanisms.

The 10th and 15th ranked genes were *GREB1L* and *GRB10*, and distinct AS patterns are suggested by the difference in mapped read counts between NSCs and NCs, especially for the intron region (Fig.7(c),(d)). In *GREB1L*, several reads mapped to the 5*’* intron region (left side of the heatmap in Fig.7(c)), and the long isoform appears to be expressed in NSCs. Our NMF-based algorithm detected such overlooked differences 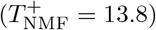 in contrast to the annotation-based approach 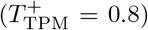. Since RamDA-seq detects not only mature mRNAs but also pre-mRNAs, many reads mapped to intron regions are considered to be derived from pre-mRNA expression [27]. Because the annotation-based algorithm does not usually use intron-mapped reads, our proposed algorithm that utilizes such information is effective for AS pattern identification, especially for genes with alternative TSSs.

For *GRB10*, numerous reads were mapped to its 5*’* intron, and cell-type-specific TSS switching likely occurs for this gene (Fig.7(d)). *GRB10* is an imprinted gene and is known for its unique TSS switch mechanism in mouse [35]. In the differentiation of mESCs into motor neurons, the expression of *Grb10* changes from the maternal to paternal allele. The upstream promoter is used for maternal expression, and the downstream alternative promoter is used for paternal expression. Therefore, the 5*’* intron-mapped reads, which are detected in only NSCs, support the alternative TSS based on the above mechanism and reflect DE patterns, observable by utilizing intron reads.

The 17th ranked gene was *PTPRN2*, and there appears to be a short unannotated transcript in NSCs (Fig.7(e)). Notably, in mouse, an alternative promoter exists downstream of *Ptprn2*, and the transcription from the promoter drives the miR-153 precursor transcript embedded in the *Ptprn2* gene region [36]. Moreover, miR-153 is highly expressed in mouse neural stem/progenitor cells (NSPCs), and the repression of miR-153 leads to differentiation, and hence, miR-153 modulates NSPCs [37]. Human miR-153 is located in *PTPRN2* [38], and therefore, the short transcript in the 3*’* region is likely a key factor that distinguishes human NSCs and NCs but is overlooked by annotation-based analysis.

The 18th ranked gene was *GPI*, and numerous reads from NCs were mapped to its central intron region (Fig.7(f)). In *GPI*, the existence and conservation of a minisatellite in its intron have been reported [39]. Although the increase in such reads might be an artifact caused by repetitive sequences, a NC-specific transcript might exist in the region.

## Discussion

In this research, we developed a novel computational approach for differential expression analysis of scRNA-seq data based on matrix factorization of mapped count data to discover overlooked DE gene regions. Matrix factorization methods, such as principal component analysis, are a practical approach to extract essential structures and uncover biological knowledge from large-scale biological data [40]. To take advantage of the large number of cells assayed in scRNA-seq data, we proposed an NMF-based approach to extract reproducible patterns and quantify differences in these patterns among groups. In particular, we used non-negative constraint to quantify DE patterns while preserving information about the group in which the patterns were expressed, and we developed a score that identifies ODEGRs by using positive maximum and negative minimum values. Such computational approaches which utilize numerical constraints based on the biological subjects can facilitate further omics studies.

We applied our algorithm to two scRNA-seq datasets and discovered several unannotated DE patterns, including DE antisense transcripts. In addition, our algorithm utilized mapping patterns in intron regions to discover overlooked alternative TSS patterns. Specifically, we detected an unannotated transcript which is a key factor for regulating differentiation. Thus, our approach has the potential to identify essential overlooked DE genes.

Although our algorithm was able to identify several intriguing ODEGRs, it remains difficult to distinguish the cause of DE transcripts such as those associated with antisense transcripts or the long unannotated transcripts of adjacent genes. In addition, the detected ODEGRs are few, and thus the impact on whole expression analyses is quantitatively small. However, our approach can discover novel transcripts and will enable further experimental and computational analyses of these transcripts, which will deepen the current understanding of the complex gene expression landscape. As shown in the validation of alternative isoform expression, our algorithm overlooked several genes with alternative isoform expression. One limitation of our algorithm is that its detection of changes involves small exons, because small changes have little effect on the objective function and are overlooked in matrix factorization. In addition, we used the count data with a 100-bp bin size (see the Methods section), which also overlooked the differences in small exons. Although this problem might be solved by using smaller bin sizes, NMF computational time and data size increase substantially with increases in matrix size, so additional improvements are therefore necessary. Moreover, our algorithm overlooks DE patterns in the filtered regions such as those with gene overlap or those with low mappability. Therefore, other approaches, such as methods based on exon–exon junction reads [17, 18], will be useful to make up for each other’s weak points and to complement annotation-based analyses.

Several effective computational expression analysis methods for scRNA-seq data, such as for cell typing and for reconstructing differentiation trajectories, have been developed so far. In this research, we have proposed a novel application of scRNA-seq data for discovering overlooked DE transcripts. Here, we have developed an algorithm for differential expression analysis between two groups, and this approach might be useful for analyzing cellular heterogeneity and discovering transcripts with an overlooked multimodal distribution.

## Conclusions

In summary, we have developed an algorithm to discover overlooked DE gene regions from scRNA-seq data. First, we confirmed that our algorithm could detect complex DE patterns such as simulated local differential expression and alternative isoform expression. Then, we applied our algorithm to two single-cell full-length total RNA-seq datasets and discovered intriguing examples of differential expression, including a transcript related to the modulation of NSPC differentiation. Our approach complements annotation-based analysis and is an effective approach for better understanding cellular regulatory mechanisms using singlecell studies.

## Methods

### Data processing

The mouse ES-PrE dataset was derived from our previous work [27], and we regarded cells 0 h and 72 h after induction as ES and PrE cells, respectively. The scRNA-seq reads were aligned to the mouse mm10 genome using HISAT2 [41] with the parameters “–dtacufflinks -p 4 -k 5 -X 800 –sp 1000,1000,” and uniquely mapped reads were selected using the BAMtools “filter” command with the parameters “-isMapped true -tag NH:1” and the SAMTools “view” command with the parameter “-q 40.” The genome-wide coverage data were generated from these mapped data using the “bamCoverage” command in deepTools(2.7.10) [42] with the parameters “–binSize 1 –smoothLength 1 – normalizeUsingRPKM.” We also quantified transcript-level expression data (i.e., TPM matrix) from scRNAseq data using the Sailfish(v0.9.2) [43] “quant” command with the parameter “-l U” and GENCODE vM9 annotation.

The human NSC-NC dataset was measured using RamDA-seq for cell populations derived from NSCs differentiated from iPS cells. The scRNA-seq reads were aligned to the human hg38 genome with STAR(v2.5.2a) [44], and the coverage data was constructed with “bamCoverage” command as mentioned above. We also quantified the transcript-level expression data (TPM matrix) with Sailfish(v0.10.0) based on GENCODE v24 gene annotation. Based on the known marker gene expression, we identified subpopulations in the data (Additional file 1: Fig. S1). In particular, we found that a subpopulation expressed some stemness marker genes, such as *SOX2, LIN28*, and *POU5F1*, and another subpopulation expressed neural marker genes, such as *ASCL1*. We regarded the cell types corresponding to those two subpopulations as NSCs and NCs, respectively.

For both datasets, we generated a mapping count data matrix for each gene region as follows. First, we extracted the transcript list so that the mean expression of a transcript *t* is over a set threshold (i.e., ∑_*c*_ log_10_(TPM_*t,c*_ + 1)/*C* > 0.5, where *C* is the number of cells). Next, we constructed the unique protein-coding gene list, which corresponds to the above transcript list. Then, we selected 6,921 and 9,359 genes from each dataset and constructed a count data matrix (100-bp bins) for each gene region from the genomewide coverage data of each cells. The gene regions were defined by the genomic start location and end location of the row of the gene in the GENCODE GTF files (vM9 for the mES-PrE dataset and v24 for the hNSC-NC dataset). We filtered the bins that contained various genes because the target gene might falsely be regarded as occurring in an ODEGR owing to the differential expression of other overlapping genes. We also filtered the bins that were derived from regions with low mappability. This is because such bins might falsely be regarded as a differentially expressed region owing to the misalignment of reads. In this research, we defined bins with low mappability as those for which the minimum of 24-bp mappability (downloaded from https://bismap.hoffmanlab.org [45]) was 0.5 or less. Then, the genes that remained with bin sizes under 100 were filtered. In this way, 4,965 and 6,491 genes were selected for differential expression analysis.

### Implementation and computational cost

We computed NMF with the *NMF* package in the R statistical computing environment [29] and used the objective function based on the Euclidean distance between the data matrix **X** and the reconstructed matrix **WH** as calculated by factorization [46]. The raw count matrix data has excessively large values in some bins, and such large values cause the underestimation of the influence of the remaining bins in the objective function. Therefore, we applied a log_10_(count + 1) transformation to the count values before NMF calculation. The scripts are available at GitHub (https://github.com/hmatsu1226/ODEGRfinder).

Since the NMF calculations of all gene regions are independent from each other, we performed NMF for each gene region in parallel using Sun Grid Engine. In the NMF analysis with *K* = 10 for the first 1,000 gene regions, the computational times were about 1.7 hours and 10.9 hours with maximum memory usage of about 240 Mb and 544 Mb for the mES-PrE and hNSC-NC datasets, respectively.

### Validation method

#### Simulation dataset

We constructed simulation data from the mES-PrE dataset. First, we calculated the mean of the logarithm of the coverage of a gene region 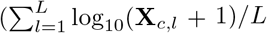, where *L* is the number of bins and *c* is the index of a cell). We then calculated the *p*-value of the *t*-test comparing this value between the ES cells and PrE cells and extracted the top 100 most significant DE genes. Second, we randomly selected a sample of count data (**X**) from these 100 DE genes, and reshaped the *C × L* matrix **X** into a *C* × *L’* matrix **X***’* (*L*’ < *L*) by averaging **X**_*c,i*_ from *i* = *l*(*b* – 1)(*L* – 1)/*L*’*J* to *b*(*L* – 1)*/L*’ for each bin *b* corresponding to 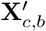. Then, we randomly selected a gene from among 4,965 genes and combined the count data for the gene using the above matrix **X***’* so that the combined matrix had the local DE pattern. However, if the two selected genes had the same DE trend, that is, both satisfied –log_10_(*p*-value) > 10 for the same side in the corresponding *t*-test, the combined matrix did not have the local DE pattern, and so we selected one of the 4,965 genes at random again. We generated a positivecontrol datasets with 1,000 datapoints as above for *L*’ = 10, 50, and 100, and we regarded the raw data as the negative-control set.

#### Alternative isoform expression definition

We defined genes with alternative isoform expression based on the TPM matrix. We defined a gene that satisfied *-* log_10_(*p*-value) for a corresponding *t*-test for 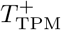 and 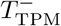 over *α* as belonging to the positive-control set, and the remaining genes as belonging to the negative-control set. We used *α* = 5, 10, and 15 and the number of genes in the positive-control set were 75, 25, and 8 for the mES-PrE dataset and 333, 95, and 51 for the hNSC-NC dataset, respectively.

### Discovery of ODEGR

We investigated the ODEGRs based on their ranked Δ*T*_NMF–TPM_ values in descending order. Even if Δ*T*_NMF–TPM_ is large, the annotation-based approach also detects the DE when *T*_TPM_ is sufficiently large. Therefore, we used 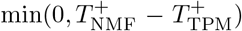 instead of 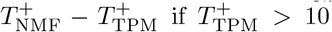 and 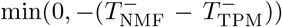 instead of 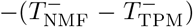 if 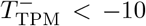 for calculating Δ*T*_NMF–TPM_ to discover overlooked DE gene regions (see Algorithm 1).

#### Algorithm 1

Calculate Δ*T*_NMF–TPM_

Δ*T* ← –*∞*

**if** 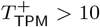 **then**

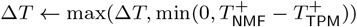

**else**

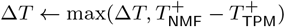

**end if
if** 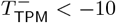 **then**

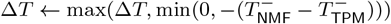

**else**

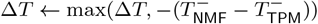

**end if**

return Δ*T*

We also considered the reproducibility of NMF results. As there is no global optimization algorithm for NMF, the result depends on the initialization. Accordingly, we calculated Δ*T*_NMF–TPM_ for a gene with three initial values generated by different random seeds, and we used only the minimum value of Δ*T*_NMF–TPM_ among the three trials to filter unreliable differences. The reproducibility of NMF and filtered genes is described in Additional file 1.

## Ethics approval and consent to participate

The use of human iPSCs derived NSPCs was approved by ethics committees at Keio University School of Medicine (admission numbers; 20130146).

## Consent for publication

Not applicable.

## Availability of data and materials

The sequencing data of hNSC-NC dataset can be accessed at the Gene Expression Omnibus under accession code GSE125288. The processed mES-PrE dataset and hNSC-NC dataset are available at https://doi.org/10.6084/m9.figshare.7410509.v1 and https://doi.org/10.6084/m9.figshare.7410512.v1, respectively. The software ODEGRfinder is available at GitHub https://github.com/hmatsu1226/ODEGRfinder.

## Authors’ contributions

HM, TH, and IN designed the study. HM designed and implemented the algorithm. HM and HOZ analyzed the mES-PrE dataset, and HM and KT analyzed the hNSC-NC dataset. TH, MU, TI, MN, and HOK performed experiments for the hNSC-NC dataset. HM, TH, HOZ, KT, and IN wrote the manuscript. All authors read and approved the final manuscript.

## Acknowledgements

We are grateful to all members of the Laboratory for Bioinformatics Research, RIKEN Center for Biosystems Dynamics Research, for providing helpful advice. We would especially like to thank Mr. Akihiro Matsushima and Mr. Manabu Ishii for their assistance with the IT infrastructure for the data analysis. We are also grateful to Jun Kohyama (Department of Physiology, Keio University School of Medicine). We would also like to thank the Center for iPS Cell Research and Application, Kyoto University, for the human iPSC clones.

## Figures Additional Files

Additional file 1 — Additional text and figures

Additional text and figures referred in the manuscript (PDF).

