## Additional file 1 for "A NMF-based approach to discover overlooked differentially expressed gene regions from single-cell RNA-seq data"

February 7, 2019

### 1 hNSC-NC dataset

We performed single-cell RNA sequencing (scRNA-seq) for a neural stem cell (NSC) population derived from human induced pluripotent stem (iPS) cells. The population was heterogeneous as shown in Fig.S1. We clustered cells into subgroups and defined cell types based on the expression of marker genes, which resulted in the NSC subgroup (red), neural cell (NC) subgroup (yellow), and Niche cell subgroup (green). We also identified some uninterpretable subgroups (pink, cyan, and purple) that are thought to be experimental artifacts. In this research, we investigated differential expression between 515 NSCs and 80 NCs.

### 2 Analysis of NMF- and TPM-based scores

We compared the NMF-based DE score $T^{+,-}_{\mathrm{NMF}}$ and the TPM based-DE score $T^{+,-}_{\mathrm{TPM}}$ for each dataset (Fig.S2(a),(b)). In brief, $T^{+,-}_{\mathrm{NMF}}$ and $T^{+,-}_{\mathrm{TPM}}$ are consistent and strongly correlated (Pearson's correlation coefficients between the two values were about 0.83 and 0.84, respectively), but a small number of genes had significantly different scores. When the absolute values $T^{+}_{\mathrm{NMF}}$ (or $T^{-}_{\mathrm{NMF}}$) and $T^{+}_{\mathrm{TPM}}$ (or $T^{-}_{\mathrm{TPM}}$) are large, the difference $T^{+}_{\mathrm{NMF}} - T^{+}_{\mathrm{TPM}}$ (or $-T^{-}_{\mathrm{NMF}} + T^{-}_{\mathrm{TPM}}$) tends to be large in several genes, which results in large $\Delta T_{\mathrm{NMF-TPM}}$ values. However, such genes are regarded as DE genes according to both scores, and their examination is contrary to our purpose of discovering overlooked DE gene regions. Therefore, we did not evaluate such differential expression in the course of describing the discovering ODEGR in the Method section of the main text.

The values of $\Delta T_{\mathrm{NMF-TPM}}$ in descending order for each dataset are shown in Fig.S2(c),(d). A small fraction of genes show large positive and negative values of $\Delta T_{\mathrm{NMF-TPM}}$, and the ODEGRs are thought to be included in the genes with large positive $\Delta T_{\mathrm{NMF-TPM}}$ values.

Our NMF-based approach tends to overlook exons with small changes owing to their small effect in the objective function and cannot detect differential expression in the filtered regions as mentioned in the main text, which results in large negative $\Delta T_{\mathrm{NMF-TPM}}$ in some genes.

### 3 Discovery of ODEGRs

#### 3.1 Reproducibility of NMF-based scores and examples of false-positive cases

There is no global minimization algorithm for NMF, and the results of NMF vary depend on the initialization. To detect reliable ODEGRs, we calculated $\Delta T_{\mathrm{NMF-TPM}}$ three times (described as trials "A", "B", and "C") with different random seeds. The overlap of the top 20 ranking genes are visualized with Venn diagrams (Fig.S3(a)(b)). Over half of the genes were common among the three trials, and there were several intriguing patterns, as mentioned in the main text. However, there were some non-overlapping genes, and

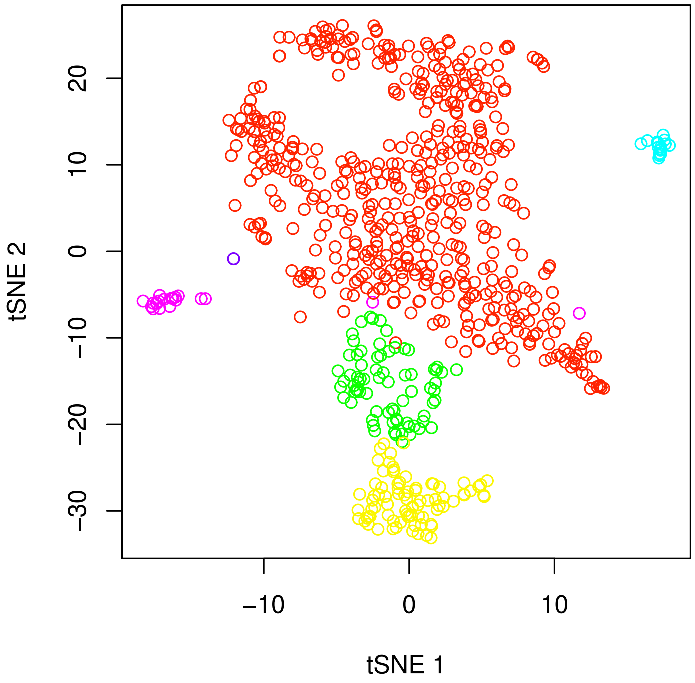


Figure S1: The t-distributed stochastic neighbor embedding analysis of the iPS cell derived neural stem cell population, colored by cluster assignment.

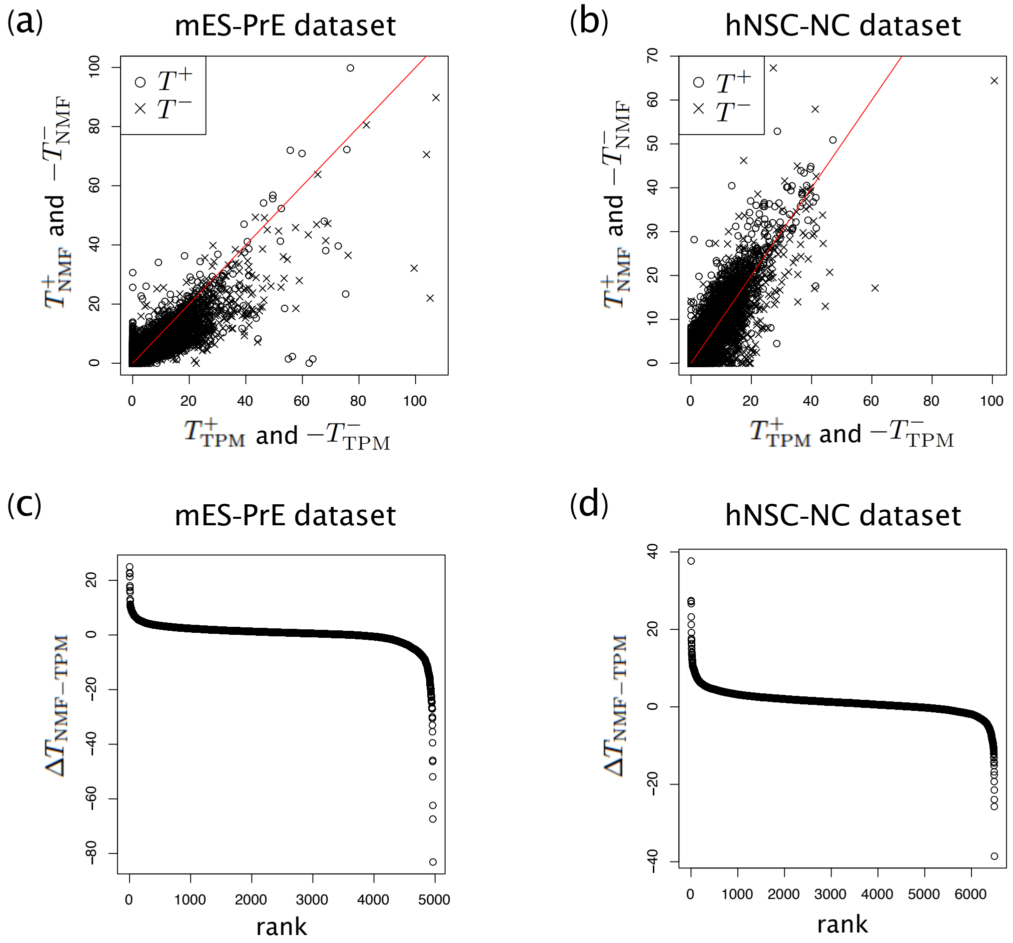


Figure S2: Comparison of $T^+_{\rm TPM}$ (or $T^-_{\rm TPM}$) and $T^+_{\rm NMF}$ (or $T^-_{\rm NMF}$) in (a) the mES-PrE dataset and (b) the hNSC-NC dataset. $\Delta T_{\rm NMF-TPM}$ values in rank order in (c) the mES-PrE dataset and (d) the hNSC-NC dataset.

we investigated those that were detected in only trial "A" for the mES-PrE dataset (Fig.S3(c)(d)). In trial "A", $T^{+}_{\mathrm{TPM}} = 0.0$ and $T^{+}_{\mathrm{NMF}} = 13.9$ for *Gfpt1*, and $T^{-}_{\mathrm{TPM}} = 0.0$ and $T^{-}_{\mathrm{NMF}} = -13.4$ for *Hspa4*. Although the NMF-based scores are high, there are no clear directional DE patterns expected from the NMF-based scores, and these two genes from trial "A" are suggested to be false-positive cases. This result happens when the NMF separates the coverage data excessively, and insignificant differences are emphasized. Such false-positive cases are usually filtered using the minimum value from multiple trials.

In the hNSC-NC dataset, most of the highly ranked genes were common to all three trials. However, there are some false-positive cases as mentioned above for the common genes. The examples of such false-positive genes are *PDZD4* and *MAPRE2*, which are ranked 3rd and 7th, respectively. These cases cannot completely be filtered even if we had used the minimum value of $\Delta T_{\mathrm{NMF-TPM}}$ from multiple trials, and further improvements are necessary to reduce such false-positive results.

### 3.2 Examples of overlooked DE patterns included in the current annotation

As mentioned in the main text, we used the GENCODE vM9 annotation, as described in our previous work, to analyze the mES-PrE dataset. The overlooked DE transcripts based on vM9 might be included in the current transcript annotations. Therefore, we reanalyzed the annotation-based approach with the current GENCODE vM18 annotation. Thus, the DE transcripts of some ODEGRs based on vM9 were detected using the current vM18 annotation. The first example is *Utrn*, which ranked 2nd, and the $T^{+,-}_{\mathrm{TPM}}$ values based on the vM18 annotation were 46.2 and -15.5, which were significantly higher and lower, respectively, than those based on the vM9 annotation ($T^{+,-}_{\mathrm{TPM}}$ values for vM9 were 0.0 and -4.4, respectively) (Figure S4(a),(b)). The second example is *Arid5b*, which ranked 7th, and the $T^{+,-}_{\mathrm{TPM}}$ values based on the vM18 annotation were 13.2 and -11.5, which were significantly higher and lower, respectively, than those based on the vM9 annotation ($T^{+,-}_{\mathrm{TPM}}$ values for vM9 were 1.1 and -0.0) (Figure S4(c),(d)). The third example is *Macf1*, which ranked 5th and corresponded to a region to which numerous PrE cell reads were mapped, and the region is annotated as exon in only the vM18 annotation (Figure S4(e),(f)).

These results show the validity of the high-ranking results and the potential of our algorithm to discover unannotated alternative splicing events.

### 3.3 Examples of overlooked DE patterns from unannotated long transcripts of adjacent genes

Some ODEGRs corresponding to high-ranking genes suggest the existence of previously unannotated long transcripts of adjacent genes, as mentioned in the main text. The key example in the mES-PrE dataset is *Echdc2*, which ranked 3rd, and the key example in the hNSC-NC dataset is *PSMB7*, which ranked 2nd. The mapping count data and annotation are visualized, including their adjacent gene regions, in Figure S5. For both genes, several reads were mapped to the 3′-end intron regions, to which only transcripts of the example genes correspond. Therefore, the reads are apparently derived from transcripts containing the retained intron of the genes. However, the mapped coverage continued from the adjacent genes, and the coverage of the 3′-end intron regions are correlated with the coverage of adjacent genes. Therefore, it is reasonable to conclude that such reads are derived from unannotated long transcripts of adjacent genes.

Such unannotated long transcripts have potential roles in gene regulation and tissue-specific imprinting, as occurs with *Copg2* and the long transcript of *MestXL* as mentioned in the main text. Thus, the detection of such phenomena is an important subject in differential expression analysis.

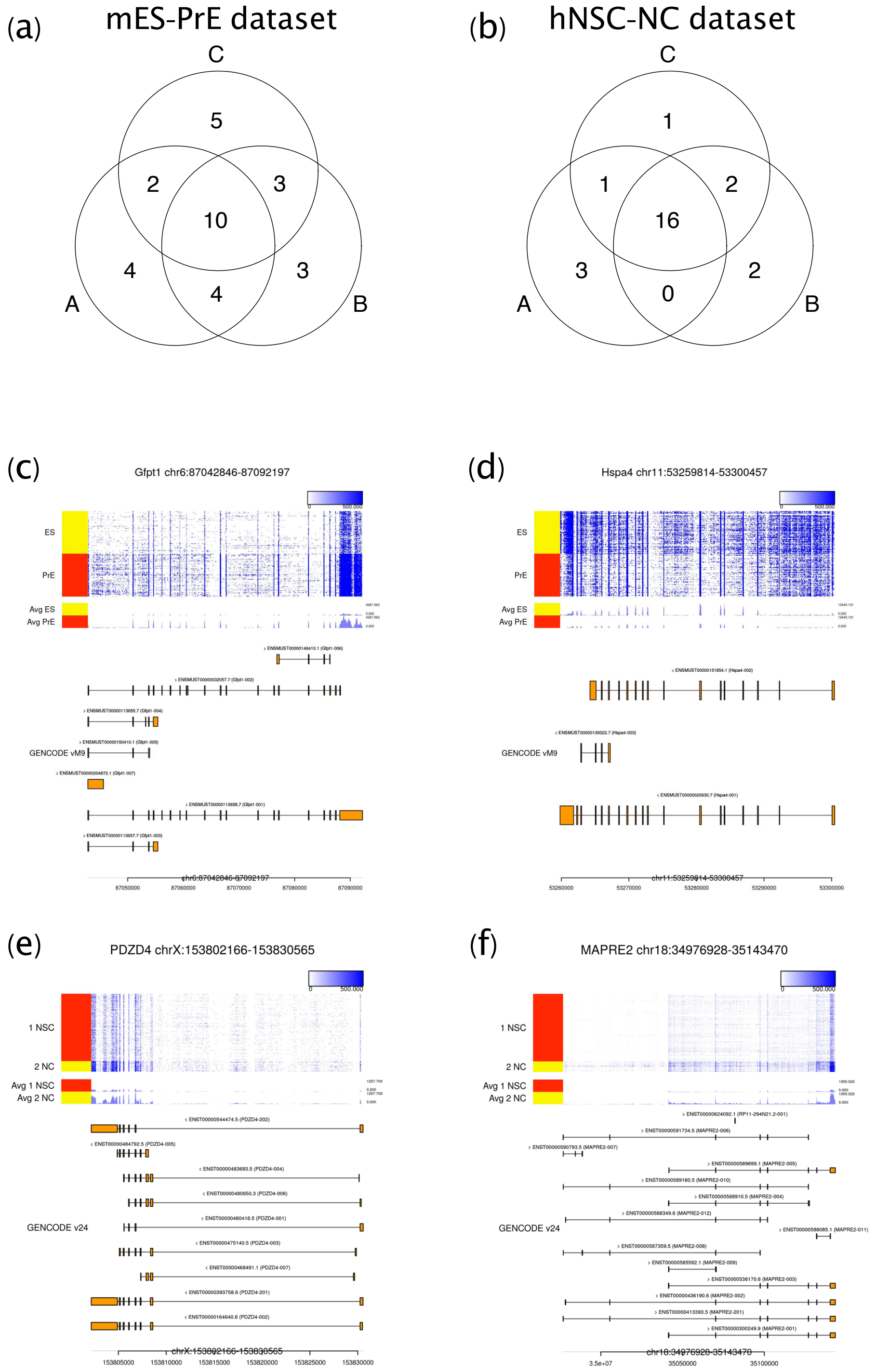


Figure S3: Venn diagrams of the top 20 ranked genes by $\Delta T_{\mathrm{NMF-TPM}}$ values in three trials with different initialization seeds for (a) the mES-PrE dataset and (b) the hNSC-NC dataset. (c) and (d) Examples of genes detected in only trial "A" in the mES-PrE dataset. (e) and (f) False-positive examples in the hNSC-NC dataset.

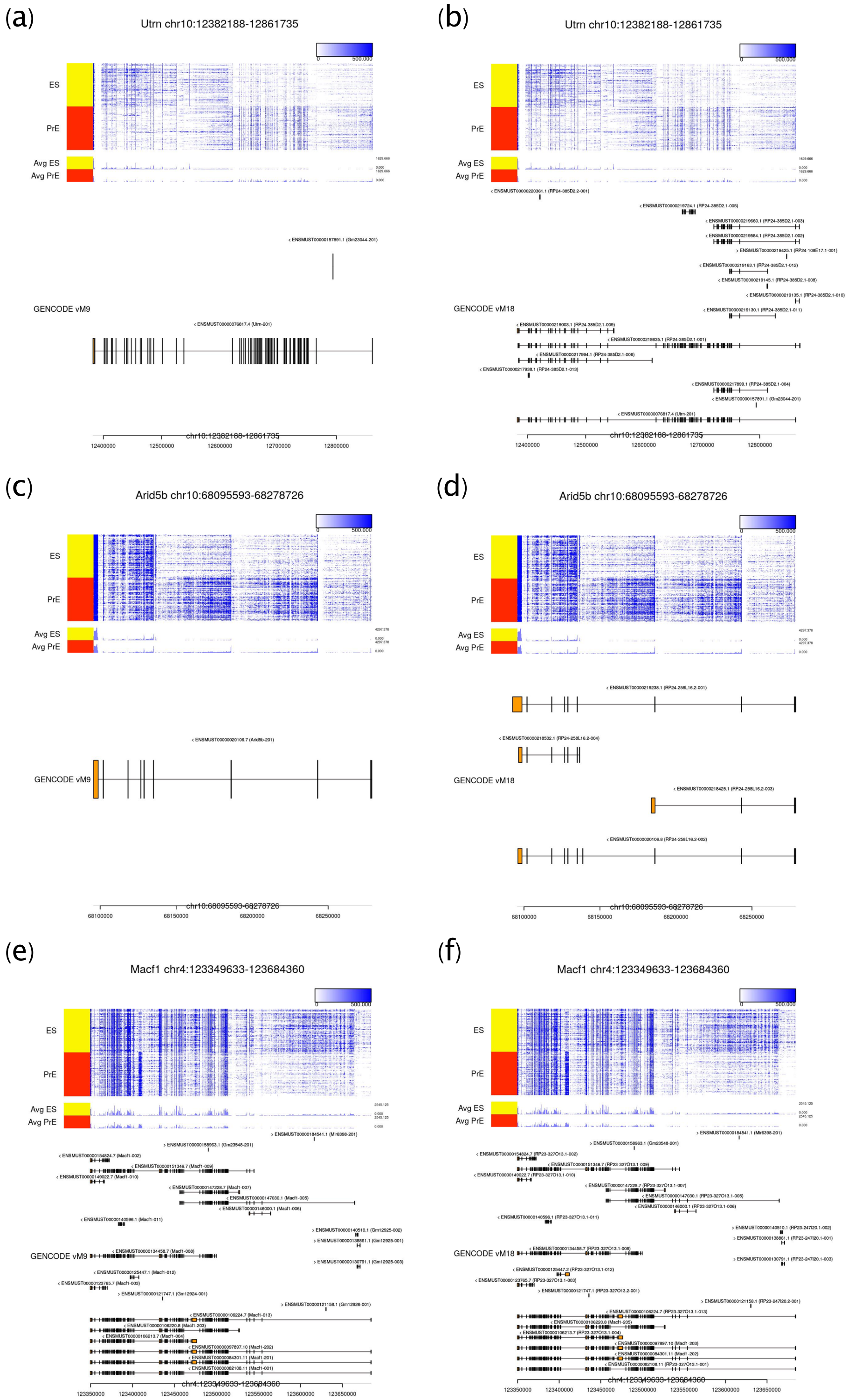


Figure S4: Examples of unexpected patterns included in the current annotation. (a) and (b) are the result of *Utrn*; (c) and (d) are the result of *Arid5b*. The left-side plots (a) and (c) correspond to the GENCODE vM9 annotation, and the right-side plots (b) and (d) correspond to the GENCODE vM18 annotation.

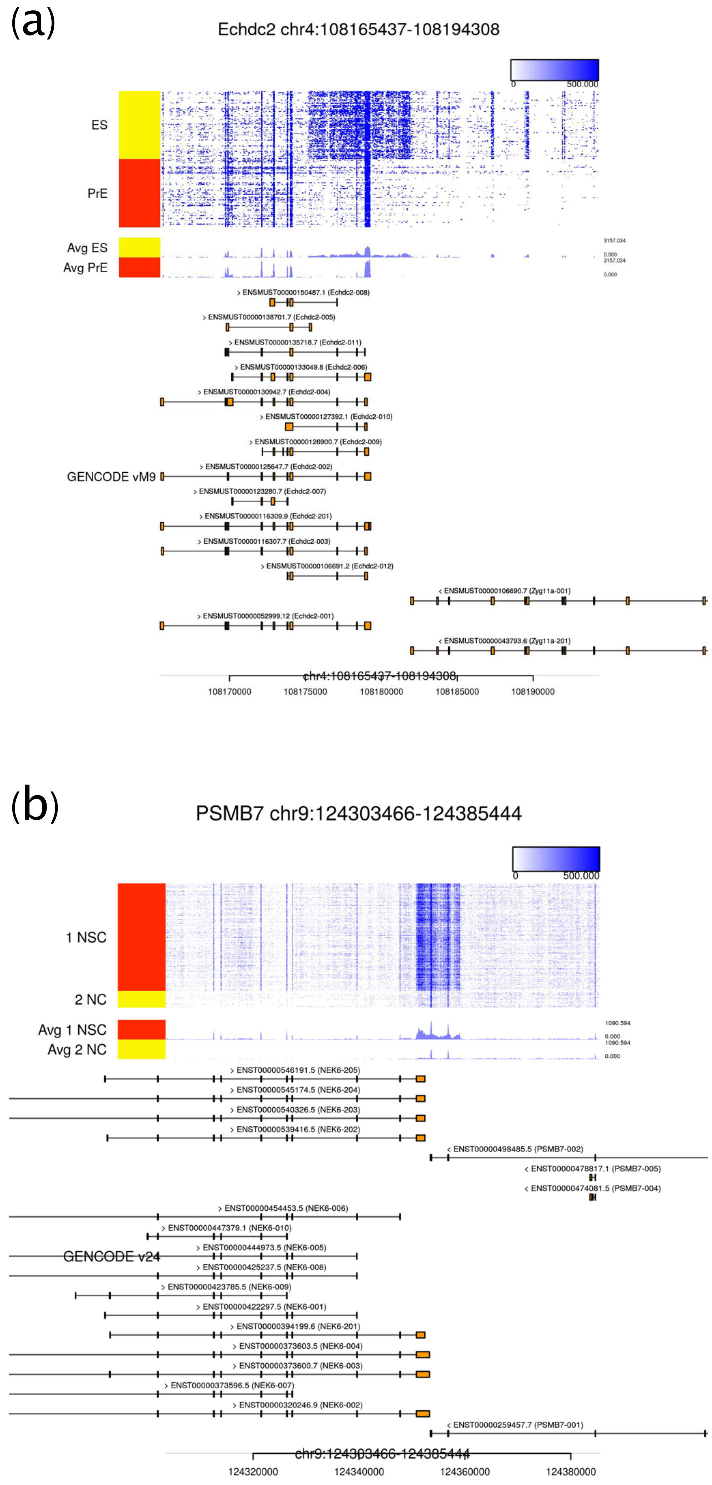


Figure S5: Examples of unexpected patterns derived from unannotated long isoforms of adjacent genes. (a) is the result of *Echdc2* in the mES-PrE dataset, and (b) is the result of *PSMB7* in the hNSC-NC dataset. We extended the visualized regions so that mapping patterns of adjacent genes are included.
